# A method to quantify autonomic nervous system function in healthy, able-bodied individuals

**DOI:** 10.1101/2020.11.24.396309

**Authors:** Shubham Debnath, Todd J. Levy, Mayer Bellehsen, Rebecca M. Schwartz, Douglas P. Barnaby, Stavros Zanos, Bruce T. Volpe, Theodoros P. Zanos

## Abstract

The autonomic nervous system (ANS), which maintains physiological homeostasis in various organ systems via parasympathetic and sympathetic branches, is altered in common diffuse and focal conditions. Sensitive, quantitative biomarkers could detect changes in ANS function, first here in healthy participants and eventually in patients displaying dysautonomia. This framework combines controlled autonomic testing with feature extraction from physiological responses. Twenty-one individuals were assessed in two morning and two afternoon sessions over two weeks. Each session included five standard clinical tests probing autonomic function: squat test, cold pressor test, diving reflex test, deep breathing, and Valsalva maneuver. Noninvasive sensors captured continuous electrocardiography, blood pressure, breathing, electrodermal activity, and pupil diameter. Heart rate, heart rate variability, mean arterial pressure, electrodermal activity, and pupil diameter responses to the perturbations were extracted, and averages across participants were computed. A template matching algorithm calculated scaling and stretching features that optimally fit the average to an individual response. These features were grouped based on test and modality to derive sympathetic and parasympathetic indices for this healthy population. A significant positive correlation (*p* = 0.000377) was found between sympathetic amplitude response and body mass index. Additionally, longer duration and larger amplitude sympathetic and longer duration parasympathetic responses occurred in afternoon testing sessions; larger amplitude parasympathetic responses occurred in morning sessions. These results demonstrate the robustness and sensitivity of an algorithmic approach to extract multimodal responses from standard tests. This novel method of quantifying ANS function can be used for early diagnosis, measurement of disease progression, or treatment evaluation.

## INTRODUCTION

The autonomic nervous systems (ANS) involuntarily regulates and integrates bodily functions derived from the physiology of internal organs like the heart, lung, spleen and intestines. That physiology also includes control over blood vessels, pupils, perspiration, and salivary glands. Regulation depends on a balance within the sympathetic and parasympathetic systems, and, it is possible to monitor real-time ANS activity by recording neural activity from candidate neural structures (Yoshimura, et al. 1994; Devor et al. 1994; Barman & Yates, 2017; Zanos, et al. 2018; Cracchiolo, et al. 2019; Masi, et al. 2019; Zanos, 2019). However, this daunting recording task would require implanted electrodes, a challenging prospect for animal experiments, no less clinical diagnosis and treatment. An alternative to direct invasive implant recording can be direct measurement of ANS-dependent physiological signals. These classes of measurements are now possible through advances in accepted, noninvasive clinical testing (Weimer, 2010). Specifically, standard techniques of autonomic testing include measuring heart rate and blood pressure during deep breathing (Shields, 2009; Coote & Chauhan, 2016; Russo et al., 2017), posture or tilt table (Porta et al., 2007; Scheen & Philips, 2012), cold pressor (Heath & Downey, 1990; Allen et al., 1992; Wirch, et al., 2006; Mourot et al. 2009; Doytchinova et al., 2016), diving reflex (Hilz et al., 1999, Hilz & Dütsch, 2005), and the Valsalva maneuver (Vogel et al., 2005; Goldstein & Cheshire, 2017; De Becker et al., 1998; Novak 2011; Gibbons et al., 2014; Doytchinova et al., 2016). Sudomotor testing relies on thermoregulatory sweat testing (TST), quantitative sudomotor axon reflex testing (QSART), sympathetic skin response (SSR), silicone impressions, the acetylcholine sweat-spot test, and quantitative direct and indirect reflex testing (QDIRT), which also belong to the standard range of provocative autonomic tests (Sumner et al., 2003; Pittenger et al., 2005; Low et al., 2006; Illigens & Gibbons, 2008).

The potential data acquired from this variety of tests can become sizeable and diverse and could exceed standard statistical evaluation. One early study developed a composite scoring scale that was able to detect generalized autonomic failure, but it could not be characterized further by disorder or for diagnosis (Low, 1993). Much attention has focused on measuring heart rate variability (HRV) as a proxy for vagal tone (Akselrod et al., 1981; Pagani et al., 1984; Beckers et al., 2001; Ducla-Soares et al., 2007; Mainardi, 2009), but no direct evidence of this relationship has been reported (Billman, 2013; Ernst 2017). HRV measures quantify fluctuations between inter-beat intervals (IBI); different time- and frequency-domain indices have been linked to short term (~five minutes) and 24-hour metrics of sympathetic or parasympathetic activity (Shaffer & Ginsberg, 2017; Stavrakis et al., 2020). Although ultra-short term (less than five minutes) HRV measures are not common, it is reported that the root mean square of successive R-R interval differences (RMSSD) is a reliable surrogate for parasympathetic activity in as low as 30 second intervals (Salahuddin et al., 2007; Baek et al., 2015; Munoz et al., 2015; Kang et al., 2016). Perhaps because HRV depends on underlying heart and respiratory rates that can be easily altered by diet, exercise, psychological stress, and medication, as well as age and gender, its reliability as a surrogate marker has remained controversial (Yamamoto et al., 1991; Perini & Veicsteinas, 2003; Young & Benton, 2018; Kyriakou, et al. 2019; Umetani et al., 1998; Stein et al., 1997; Zhang, 2007). Besides HRV, other ANS assessment techniques require drug induced response (baroreflex sensitivity), additional equipment (imaging), or invasive intraneural microelectrodes (direct muscle and sympathetic nerve activity measurement), making these assessments more difficult to administer (Goldberger et al., 2019; Stavrakis et al., 2020).

The dearth of effective, reliable, and reproducible data-driven approaches to quantify non-invasive recording modalities while administering a full battery of tests drove us to apply signal processing, machine learning models, and decoding algorithms on a set of commonly used clinical measurement of physiological signals in an attempt to derive a better understanding of autonomic function and dysregulation. Herein, we test whether this metric of autonomic function and responses is sensitive enough to identify subtle ANS deviations in health, able bodied control participants. Moreover, this approach demonstrated significant deviations of ANS responses correlated with BMI, and also showed trends related to circadian rhythm. Such noninvasive, quantitative autonomic function metrics might well enable objective measures of disease states and provide a useful tool for diagnosis and disease management.

## METHODS

### Human participants

This study recruited and enrolled 21 healthy, able-bodied participants between the ages of 18-60 years and a body mass index (BMI) less than 30. The mean age (SD) of the participants was 29.9 (6.5) years, with sixteen males and five females. The mean BMI was 24.4 (2.9). Exclusion criteria were: history of cardiac arrhythmia, coronary artery disease, autoimmune disease, chronic inflammatory disease, anemia, malignancy, depression, neurologic disease, diabetes mellitus, renal disease, dementia, psychiatric illness including active psychosis, or any other chronic medical condition, treatment with anti-cholinergic medication, current tobacco or nicotine use, smoking, pre-existing neurological disease, pregnancy, and implantable electronic devices. Participants were asked to fast and refrain from caffeine for at least four hours prior to testing. This study was approved by the Northwell Health Institutional Review Board, IRB #19-0461 and registered with Clinicaltrials.gov, identifier NCT04100486.

### Autonomic testing sessions

Testing sessions occurred in a lab office setting with no external noise or distractions. Lighting was set such that significant pupil changes were detected. Each participant attended four testing sessions: two in the morning and two in the afternoon, over the course of two weeks. Five autonomic tests were performed, with the order randomized for each participant and for each testing session (Figure 1a). In the squat test, the participant actively stood still and calm for one minute, sat in a single deep squat for one minute, and then stood again for the final minute, all in succession. For the cold pressor test, the participant immersed his or her right hand in ice water (< 5° C) for at least 30 seconds and up to three minutes; after the first 30 seconds, the participant was allowed to remove their hand at their own will. During deep breathing, the participant followed a visual cue on a dimly lit computer monitor to time their respiratory rate to six breaths per minute for seven minutes. The diving reflex test was administered by placing a refrigerated gel-filled compress on the participant’s face for one minute, followed by one minute of recovery. Lastly, the Valsalva maneuver was performed by a forceful attempted exhalation, “expelling” air while keeping the mouth and nose shut for 15 seconds, followed by one minute of recovery.

**Figure 1.**
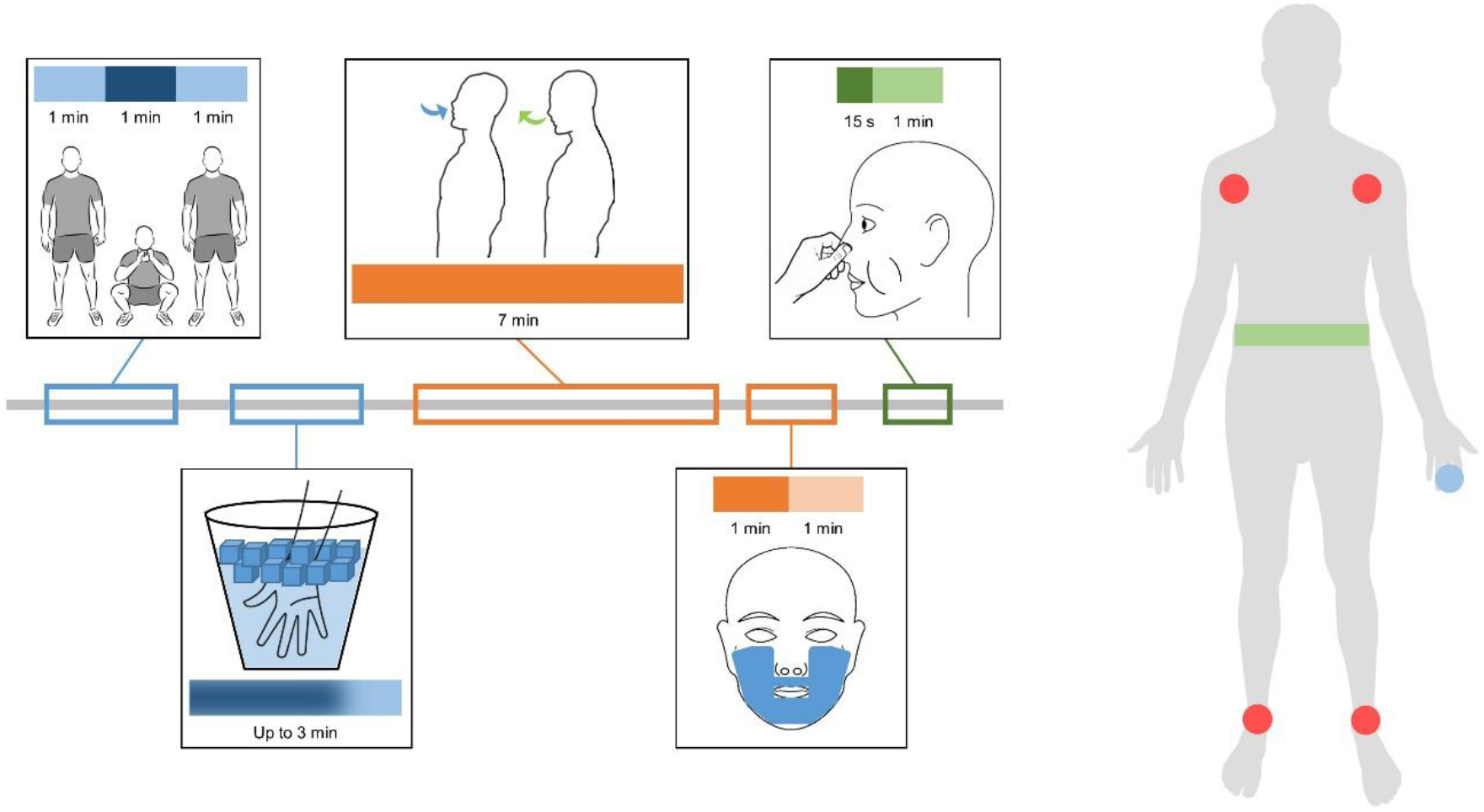
Autonomic testing and monitoring physiological signals. (a) Shown is a sample timeline of autonomic tests performed in each session. The tests include a set of sympathetic tests (standing-squatting-standing [one minute of each, in succession] and cold pressor test [immersion of hand in ice water for up to three minutes]), a set of parasympathetic tests (deep breathing [respiratory rate of six breaths per minute for seven minutes] and diving reflex test [refrigerated gel-filled compresses on the face for one minute with one minute of recovery]), and Valsalva maneuver (restricted and forced exhalation for 15 seconds with one minute of recovery). (b) Physiological signals for each participant were recorded by six lead electrocardiogram (in red, wires attached to four foam adhesive electrodes placed at each shoulder and each ankle), a respiratory belt (in green, around the torso), and noninvasive blood pressure cuff (in blue, small inflatable cuff on middle phalanx of middle finger on left hand).

### Physiological recording

In each session, the participant’s cardiovascular data were captured by noninvasive sensors (Figure 1b): six lead electrocardiography, respiration, and blood pressure. The six lead ECG wires were attached to four foam adhesive electrodes placed at each shoulder and each ankle, while the respiratory belt was wrapped and tightened around the torso. Blood pressure was recorded by a small non-invasive and inflatable cuff around the middle phalanx of the middle finger on the left hand. Additional sensors were attached to capture electrodermal activity (EDA) (dry, metal electrodes on two fingers) and pupil diameter (Tobii Pro Glasses 2, Tobii Pro AB, Stockholm, Sweden). All of the recordings were transmitted through that data acquisition system and software (LabChart, ADInstruments, Sydney, Australia) at a sampling rate of 1 kHz. All signals were marked and synchronized with experimental cues aligned with each test. Each participant’s BMI was recorded and monitored as the main characteristic to be predicted from autonomic signals. Continuous autonomic data from each participant and multiple non-invasive sensors, including ECG, blood pressure, respiration, EDA, and pupil diameter, were measured at multiple time points relative to rest and onset of the autonomic tests. Individual responses were characterized by an approach based on extracting unique features from recorded data to identify significant responses and compute physiologically relevant and plausible indices that are representative of deviations from resting ANS function; these methods were not linked with previously developed indices, tools, or features.

### Signal processing

A 60 Hz notch filter was applied to remove line noise from raw cardiovascular data signals. Pupil diameter data was calibrated based on gaze location; since illumination and perceived brightness can influence pupil size, measured pupil diameter was normalized to extract the autonomic response. Participants were not focused on any near objects, so pupil reactions due to accommodation were expected to be minimal and removed through averaging. The RGB values of the gaze location were converted to relative luminance values, and this was linearly fit to pupil diameter. By dividing by the slope of the line, the pupil sizes were normalized to account for effects of luminance.

Only modalities with a significant response to an autonomic test were considered for analysis. The mean μ and standard deviation σ were calculated for the baseline (before each autonomic test). A threshold for significance was determined as 1σ over baseline, a common way to define a threshold in signal detection theory (Merfeld, 2011); only peaks of the average response for a modality and test above this threshold were considered for analysis.

### Feature extraction

Each participant’s heart rate, mean arterial pressure, and RMSSD responses to the squat test, cold pressor test, diving reflex test, and Valsalva maneuver were characterized by a template matching method (Figure 2). The comparisons between individual responses and the average response template in each epoch were quantified to reflect the autonomic response. An individual response may be delayed or occur sooner than the average response. Individual responses can also be shorter or longer in duration and smaller or larger in amplitude than an average response. To quantify these deviations, the parameters that minimize the following objective function were estimated:

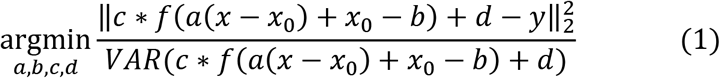

**Figure 2.**
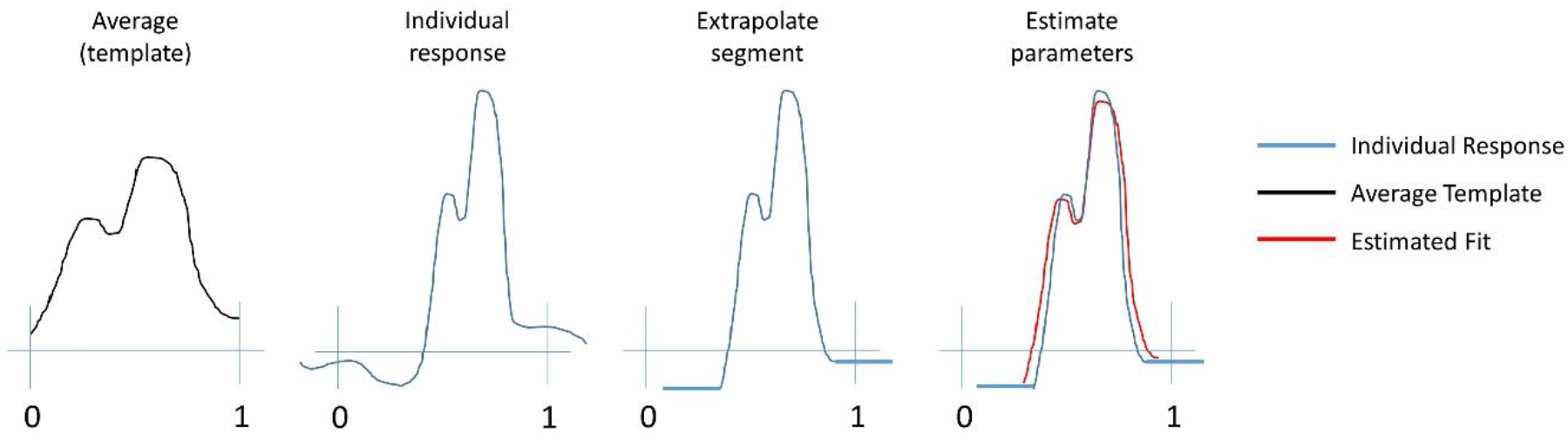
Schematic of template matching method. Shown is a schematic of the template matching method to quantify autonomic responses. The average response within an epoch (a) is the template to fit the individual response (b). Once the individual response is trimmed and extrapolated for the same epoch (c), four parameters are estimated to best fit the template to the response (d). Parameters are estimated by minimizing the normalized sum of the squared error (Equation 1). The parameters quantify how the individual response compares to the average template in duration scale and amplitude scale, as well as delay and vertical offset. The duration scale is reported as variable H and the amplitude scale and vertical offset are reported as variable V.

In equation (1), the normalized sum of the squared error is minimized. *f*(*x*) is an average response template centered at time *x_0_* and *y* is the trimmed individual response. The parameter *a* scales the duration of the response, and the parameter *b* delays or advances the response. The paramter *c* scales the amplitude of the template and will be reported as Vs, and *d* vertically offsets the template and will be reported as Vo. Since the average response of the template is normalized by the baseline period, Vs and Vo can be combined and reported as a single parameter V = Vs + Vo that describes the net scaling of the individual response with respect to the template. The parameter H is reported to represent the parameter *a* to define the duration scaling for each individual response.

For the deep breathing test, only heart rate changes were considered. The differences between maximum and minimum heart rate for each breathing cycle in the first two minutes of the test were averaged for each individual. This value was divided by the average peak-to-peak heart rate over all individuals to calculate a scale-like feature.

Once features were calculated, they were separated by the type of autonomic response: sympathetic or parasympathetic. Both branches of the autonomic nervous system are active during each autonomic test, as reflected by changes in heart rate and blood pressure. Based on literature (Victor et al., 1987; Sandroni et al., 1991; Marfella et al., 1994; Kinoshita et al., 2006; Shields, 2009) and observed data, responses were categorized by modality for each test, shown in Table 1. For example, responses that led to an increase in heart rate were considered as driven by the sympathetic nervous system, while slowed heart rates were attributed as primarily affected by the parasympathetic nervous system. In the squat test, during the phase from squat to stand, only scale (V) was used to represent sympathetic activity while only stretch (H) was used to represent parasympathetic activity (Du et al. 2005; Droguett et al., 2015). During deep breathing, the calculated scale is used to represent parasympathetic activity. For all other tests and modalities, both scale and stretch features were used in tandem as surrogates for sympathetic or parasympathetic responses.

**Table 1.**
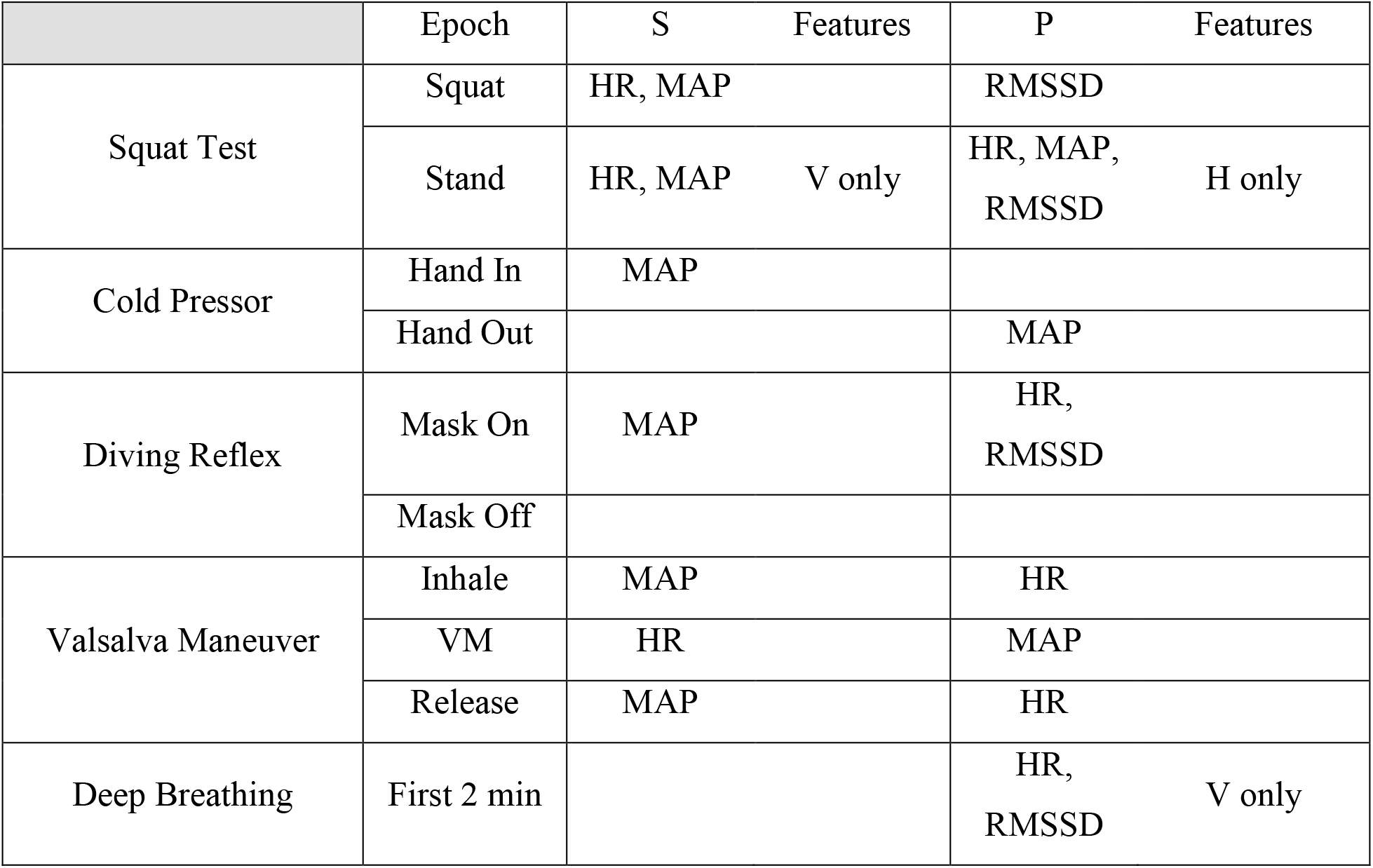
Autonomic response classification. Responses were categorized as sympathetic or parasympathetic-driven based on literature and observed data. For most responses, both scale (V) and stretch (H) features were used together for classification. In the stand epoch during the squat test, only V were classified as sympathetic and only H were classified as parasympathetic. Similarly, only a scale feature during the deep breathing test was considered as parasympathetic. Other responses were ignored, as they could not be classified as primarily sympathetic or parasympathetic branch driven.

### Modeling and validation

Classified signals were correlated with measured characteristics of the patient cohort, particularly participant BMI. Linear regression models were applied to fit the average of collected features from all modalities and testing epochs for each patient. Significant trends were validated by 10 repeats of 7-fold cross-validation; data from three patients were left out of the modeling for each fold, and samples were reshuffled for every repetition. The *p-* value was determined by a *t*-statistic; values less than 0.05 were considered significant.

## RESULTS

### Monitoring physiological signals to compute average responses

Cardiovascular, pupil dilation, and EDA signals were synchronously recorded for each participant, with raw recordings for ECG and blood pressure used to calculate heart rate, HRV (RMSSD), and mean arterial pressure (Figure 3). The individual responses of calculated signals during each test were averaged to determine a response template (Figure 4 and Figure 5). For each test and each modality, baseline-normalized responses were averaged across all participants and sessions. For HRV, we calculated the RMSSD feature for the three tests with intervals longer than 60 seconds, since this HRV measure requires 60 seconds of activity to be calculated accurately (Salahuddin et al., 2007; Baek et al., 2015); heart rate, mean arterial pressure, pupil dilation and EDA measurements were averaged for all tests. The average responses for each modality and each test at epochs with phasic changes in activity were used as temples to compare to the individual responses for each modality and test. In the squat test, the epoch corresponded with posture changes from stand to squat and squat to stand. For the cold pressor test, the epoch reflected the participant immersing their hand in the ice water, while the epoch in the diving reflex test corresponded with the cold mask being placed onto the face. The Valsalva maneuver had four epochs that reflect dynamic changes in both heart rate and blood pressure. For deep breathing, the epoch was the entire first two minutes of the task.

**Figure 3.**
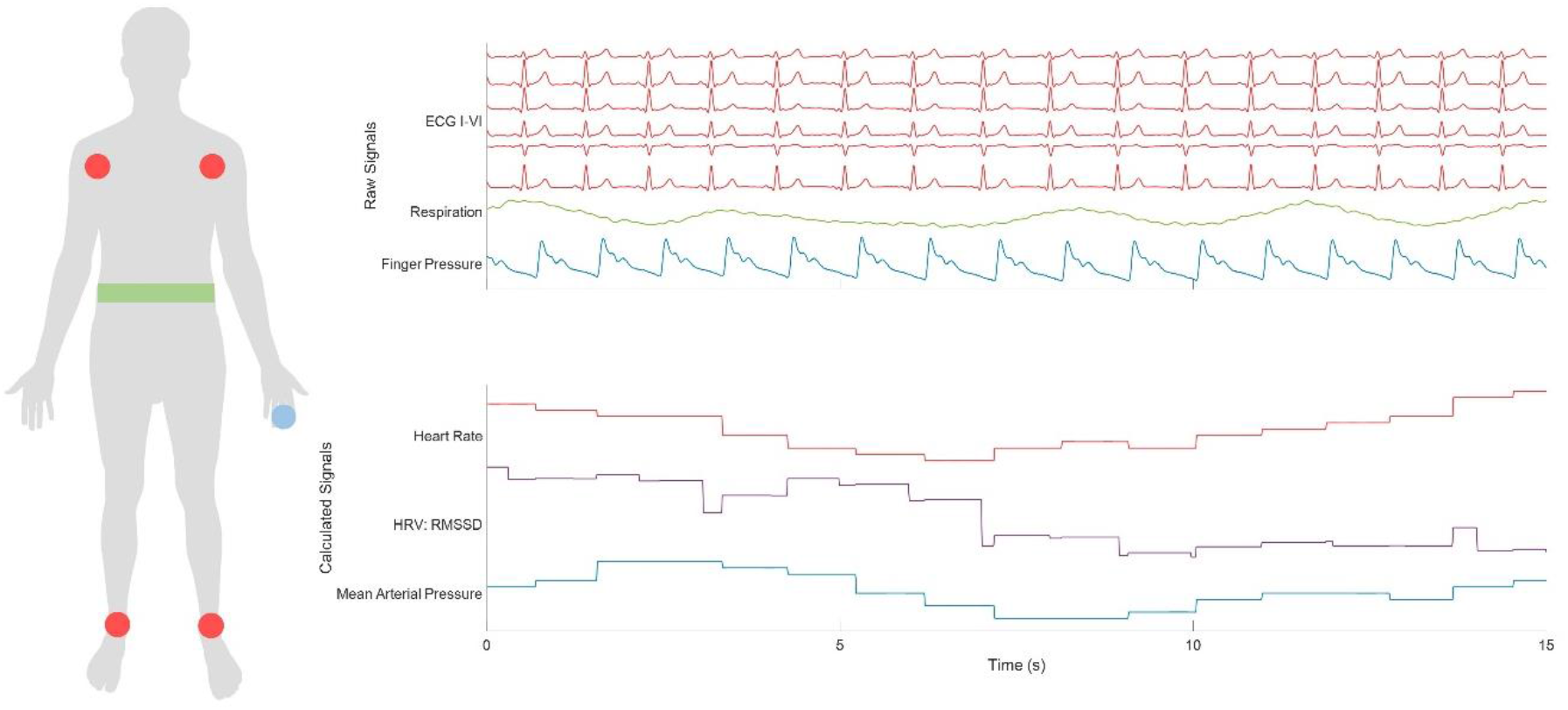
Monitoring and calculating physiological signals. (a) Physiological signals for each participant were recorded by a six lead electrocardiogram (filled red circles, wires attached to four foam adhesive electrodes placed at each shoulder and each ankle), a respiratory belt (green belt, around the torso), and noninvasive blood pressure cuff (filled blue circle, small inflatable cuff on middle phalanx of middle finger on left hand). (b) 15 seconds of raw signals from these sensors are shown in the upper panel, with six channels of ECG, one channel of respiration, and one channel of finger pressure. Derived signals in the lower panel include heart rate from the ECG, heart rate variability (RMSSD) from interbeat intervals, and mean arterial blood pressure.

**Figure 4.**
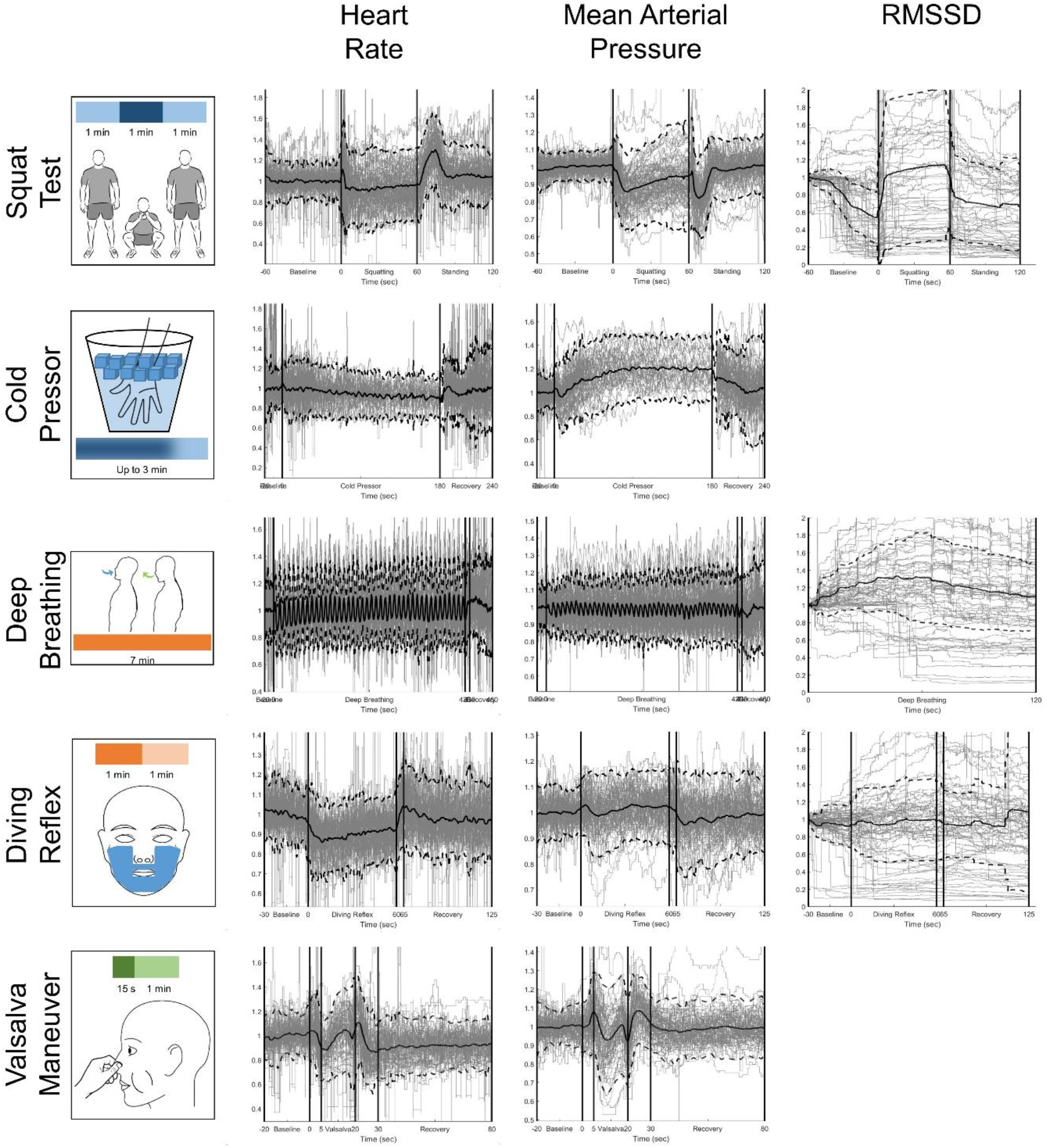
Average heart rate, mean arterial blood pressure, and RMSSD during autonomic testing (76 sessions). The individual calculated responses (gray lines) were accumulated and averaged (black line) to extract an average response for each modality during each test. Each column represents a different calculated signal (heart rate, mean arterial blood pressure, and RMSSD). The dotted black traces correspond to a 95% confidence interval. RMSSD was not calculated for the cold pressor test and Valsalva maneuver due to time constraints necessary to accurately convey heart rate variability. (a) Squat Test: vertical lines reflect changes in posture from standing to squatting and then squatting to standing. (b) Cold Pressor Test: the first vertical line reflects when the participant immersed their hand into the ice water. The second vertical line represents the maximum of three minutes. The average trace only represents the individual traces available at that time point, as participants removed their hand at their own discretion. All participants kept their hand in the ice water for at least 30 seconds. (c) Deep Breathing: vertical lines reflect when the deep breathing rate (6 breaths/minute) began and ended. In the third column for the RMSSD, only the first two minutes were analyzed. (d) Diving Reflex: vertical lines reflect when the refrigerated gel-mask was placed on and removed from the participant’s face. A five second removal period is designated before the one minute of recovery. (e) Valsalva Maneuver: vertical lines reflect the phases of the effort, from baseline, five seconds designated for inhalation and preparation, 15 seconds of the Valsalva maneuver, 10 seconds at the end of maneuver, and a final minute of recovery.

**Figure 5.**
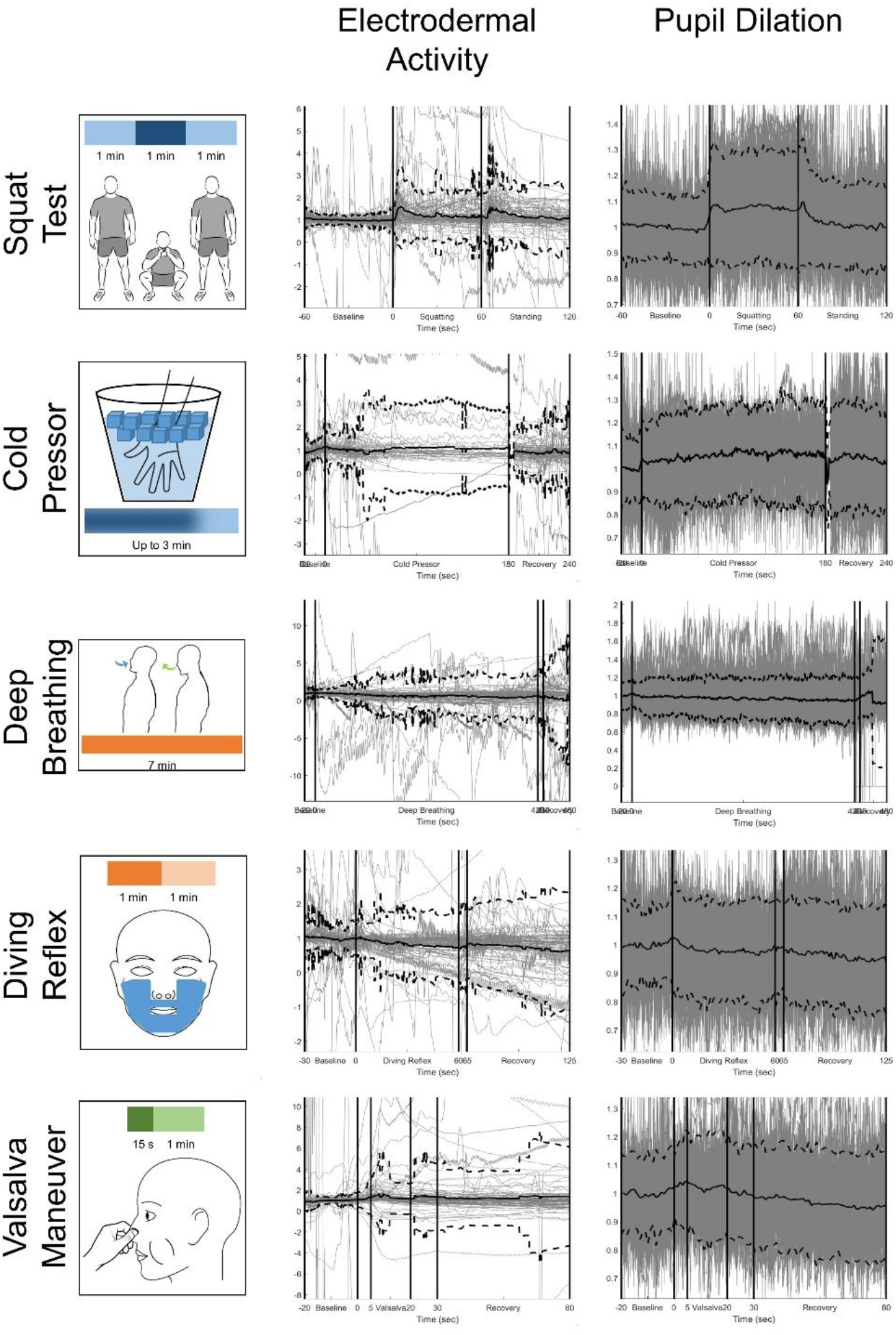
Average electrodermal activity and pupil dilation during autonomic testing (76 sessions). The individual calculated responses (gray lines) were accumulated and averaged (black line) to extract an average response for each modality during each test. Each column represents a different calculated signal (heart rate, mean arterial blood pressure, and RMSSD). The dotted black traces correspond to a 95% confidence interval. RMSSD was not calculated for the cold pressor test and Valsalva maneuver due to time constraints necessary to accurately convey heart rate variability. (a) Squat Test: vertical lines reflect changes in posture from standing to squatting and then squatting to standing. (b) Cold Pressor Test: the first vertical line reflects when the participant immersed their hand into the ice water. The second vertical line represents the maximum of three minutes. The average trace only represents the individual traces available at that time point, as participants removed their hand at their own discretion. All participants kept their hand in the ice water for at least 30 seconds. (c) Deep Breathing: vertical lines reflect when the deep breathing rate (6 breaths/minute) began and ended. In the third column for the RMSSD, only the first two minutes were analyzed. (d) Diving Reflex: vertical lines reflect when the refrigerated gel-mask was placed on and removed from the participant’s face. A five second removal period is designated before the one minute of recovery. (e) Valsalva Maneuver: vertical lines reflect the phases of the effort, from baseline, five seconds designated for inhalation and preparation, 15 seconds of the Valsalva maneuver, 10 seconds at the end of maneuver, and a final minute of recovery.

### Extracted features from individual responses

From all the physiological modalities we monitored, participants’ cardiovascular measures (HR, MAP, and RMSSD) registered average responses with a peak above 1σ of their baseline, thus deemed significant responses (as detailed in Methods), while the pupil dilation and EDA measures did not show a consistent, significant response and were discarded from further processing and feature extraction (Figure 5). Figure 6 shows examples of individual responses, the calculated features, and the corresponding transformation of the average signal based on those features. The blue traces in each panel represent an individual’s phasic response within a single epoch for an autonomic test, while the black trace in each panel is the template from the average response over all individuals and all sessions for that modality and that test. After applying the template matching algorithm, the calculated duration scaling (H), vertical scaling (Vs), and the vertical offset (Vo) can transform the average template to the red trace, which shows the estimated fit in each case. The features illustrated in these examples are representative across individuals, tests, and modalities.

**Figure 6.**
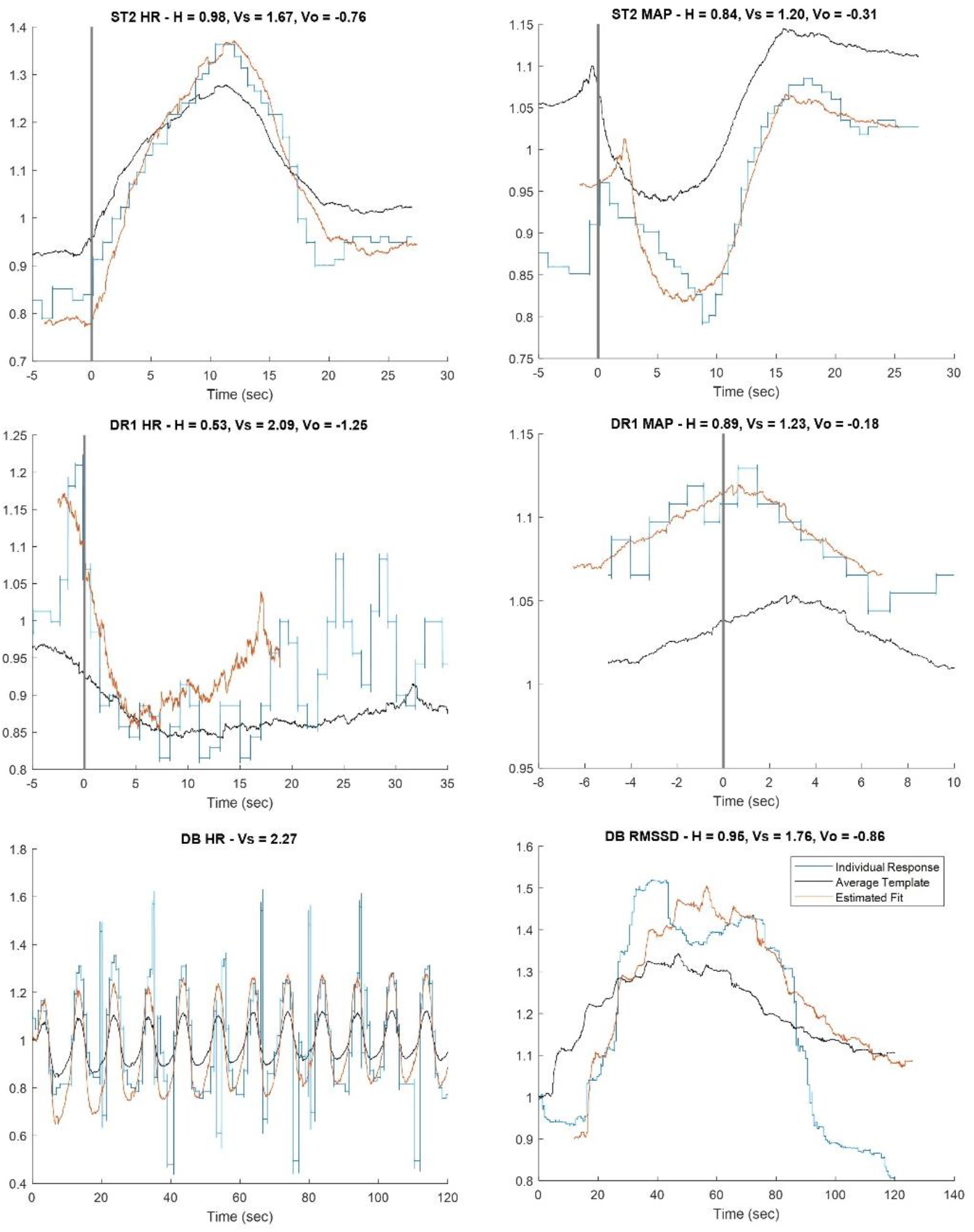
Example of individual responses and corresponding calculated features. Here are examples of individual phasic responses to autonomic testing (blue trace) and the transformed template (red trace) resulting from stretching and scaling the average template (black trace) to best match the individual response. The calculated features (in title of each panel) represent scaling amplitude (V) and duration (H). (a) Heart rate change in the transition from squatting to standing. Participant stood at t=0 (black vertical line). (b) Mean arterial pressure changes in the transition from squatting to standing. Participant stood at t=0 (black vertical line). (c) Heart rate change when putting on the refrigerated mask to initiate the diving reflex. The mask is placed at t=0 (black vertical line). (d) Mean arterial pressure change when putting on the refrigerated mask to initiate the diving reflex. The mask is placed at t=0 (black vertical line). (e) Heart rate fluctuations during first two minutes of deep breathing task. (f) Heart rate variability RMSSD calculated during the first two minutes of the deep breathing task.

Closer inspection of individual responses can provide intuition into physiological meaning of the vertical and horizontal scaling, as well as the vertical offset. As evident in the examples plotted in Figure 6a and 6b, the features extracted during the squat test reveal certain response properties, specifically in the transition from squatting to standing (at t = 0), for heart rate and mean arterial pressure, respectively. In the first example (Figure 6a), heart rate response was faster (H < 1) and larger (Vs > 1) than the average response during this squatting-standing transition. Meanwhile, in the second example (Figure 6b), the mean arterial pressure response was also faster (H < 1) and larger (Vs > 1) than the average response.

Similar interpretations can be made from the responses during the diving reflex test (Figure 6c-d), where t = 0 indicates the time that the refrigerated mask was placed on the participant’s face; 6c shows an example of a heart rate response, and 6d shows an example of a response in mean arterial pressure. In both cases, the individual’s responses were faster (H < 1) and larger (Vs > 1) than the average response.

For the deep breathing test, the vertical scaling feature of the heart rate response scales the average peak-to-peak heart rate over all individuals based on the maximum and minimum heart rates in each breathing cycle within the first two minutes of the test. The response in the specific example of Figure 6e was more than twice as large as the average response (Vs = 2.27). As mentioned in the methods, since deep breathing was visually guided, all peaks were entrained to the respiratory cycle, and there was no need for horizontal scaling feature extraction.

Similarly, the features and fit of the RMSSD measure of HRV during the first two-minute interval of deep breathing reveal the vertical and horizontal scaling using the template matching algorithm; the RMSSD response features, as calculated for the specific example in Figure 6f, reveal a faster (H < 1) and larger (Vs > 1) than the average RMSSD response.

### Sympathetic and parasympathetic features and their correlations to BMI and session time

The strength of this analysis is that all of the features from all of the tests could be interpreted as vertical and horizontal scaling measurements. The data features were grouped by vertical and horizontal scaling and by sympathetic and parasympathetic drive. Next, the features were averaged for each participant over all four testing sessions (see Table 1). These averages (and corresponding standard deviations) were correlated with physiological characteristics (BMI and age). Figure 7 represents sympathetic features (top row), and parasympathetic features (bottom row), left and right columns represent H (duration scale) and V (amplitude scale), respectively. Each data point is the average (SD) calculated feature for a single participant. A regression demonstrated a significant increase in sympathetic amplitude scaling with increasing BMI (*p* = 0.000377). Furthermore, this result was validated by 10 repeats of 7-fold cross-validation; the data left out was used to predict BMI based on the trend line produced. The mean absolute percentage error was approximately 11.7% with a standard deviation of 0.579.

**Figure 7.**
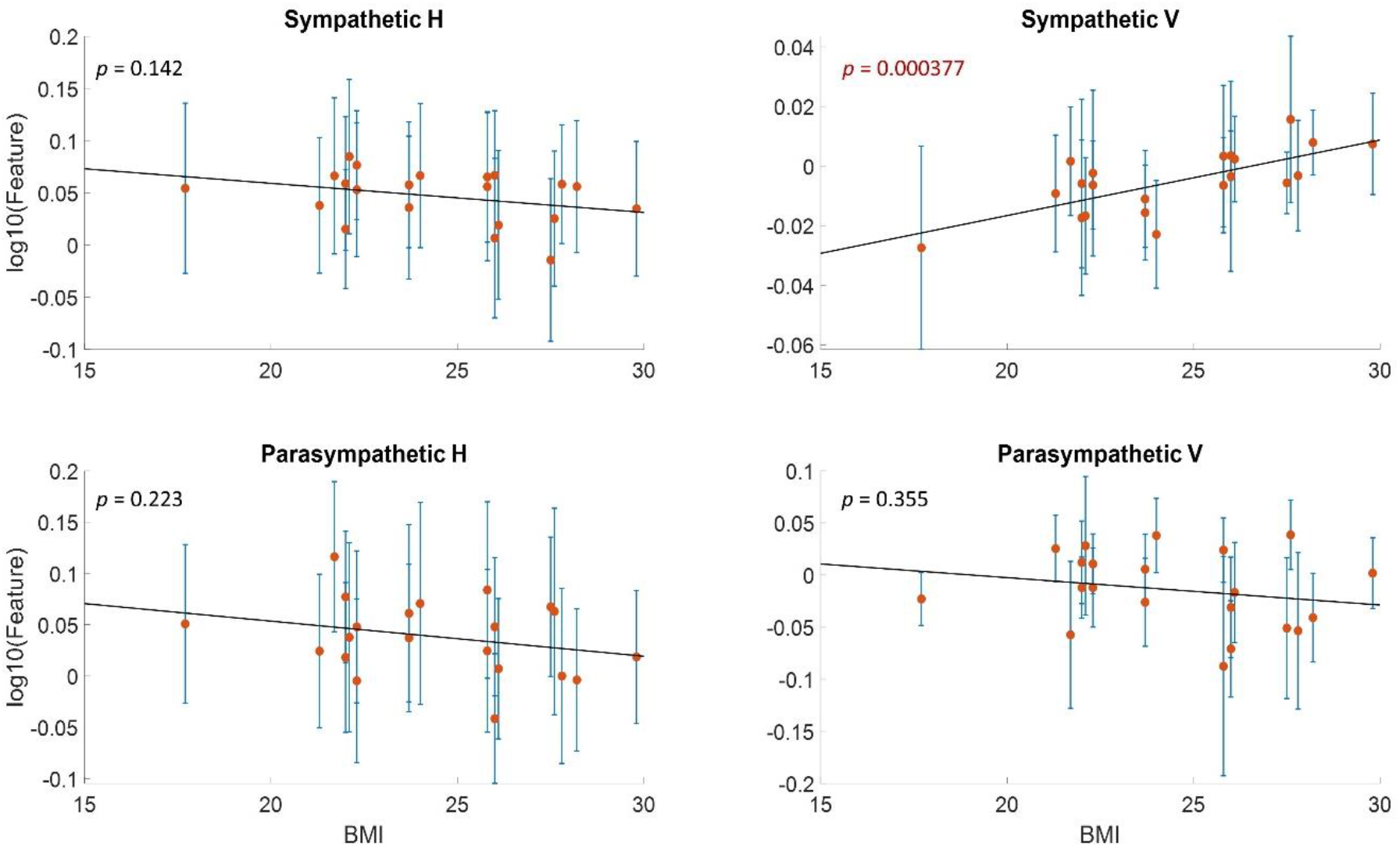
Calculated features vs BMI. Average calculated features for each participant as a function of their BMI. For each type of features (top row is sympathetic and bottom row is parasympathetic, first column represents H [duration scale] and second column represents V ([amplitude scale and offset]), each point represents the average for a single participant, while error bars show standard deviation. Line of best fit was approximated, with p-values for each subplot.

To test for an effect of circadian rhythm, the grouped features were compared across morning and afternoon sessions. The average feature for each individual was calculated over two AM sessions and two PM sessions; in Figure 8a, each trace represents a single individual, where blue and orange traces represents higher values in AM or PM sessions, respectively. These trends are summarized in Figure 8b; most (14 and 12, respectively) participants had longer (H > 1) sympathetic and parasympathetic responses in PM sessions. For vertical scaling features, larger (V > 1) sympathetic responses were observed in PM sessions for 14 participants; parasympathetic vertical scaling features were larger in AM sessions for a higher number of participants (11). Two participants did not have both morning and afternoon sessions and were left out of this analysis.

**Figure 8.**
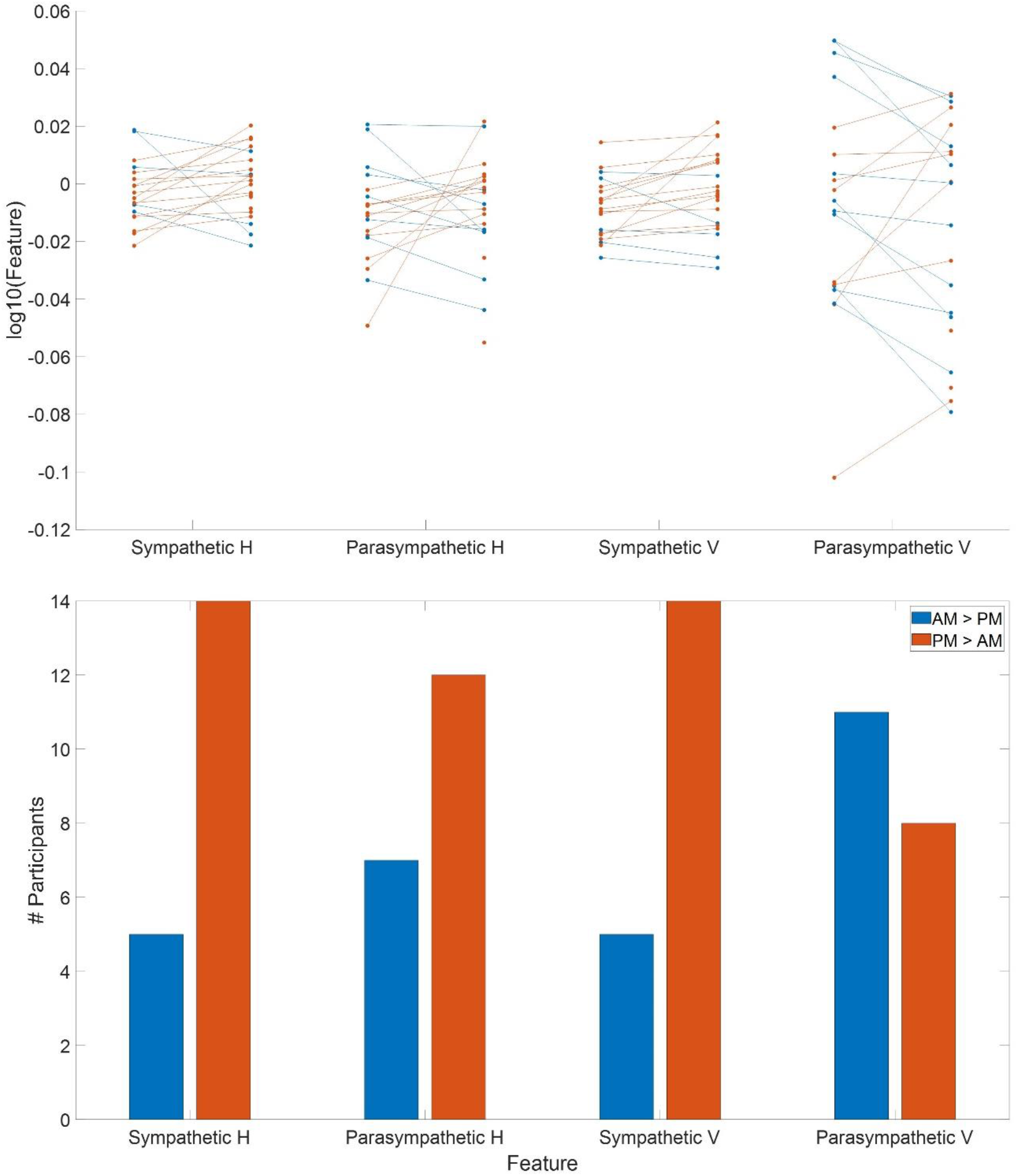
Calculated features in AM and PM. The average feature for each type for each individual was calculated for two AM sessions and two PM sessions. (a) Each trace represents one individual, where blue traces represent higher values in AM sessions and orange traces represent higher values in PM sessions. (b) Summary of the number of participants with greater feature values in either AM or PM sessions.

## DISCUSSION

Recent studies have been focused on developing quantitative standards based on biomarkers to aid with diagnosis, prognosis, and estimates of treatment efficacy (Lötsch & Ultsch, 2018). To this end, this work sought to develop a reproducible and sensitive metric to quantify significant changes in ANS function by determining features that represented duration and amplitude scales compared to the average response. Instead of a priori developed indices, a template matching method was used to estimate indices that characterized physiologically-relevant indices that represent slight deviations in sympathetic and parasympathetic individual responses. The trends revealed by these data show that sympathetic amplitude responses significantly increased and parasympathetic responses decreased with increasing BMI. These results are consistent with past work that also demonstrated increased sympathetic activity, actually significant overactivity, and decreased parasympathetic activity in patients with increased BMI greater than 25 (Molfino et al., 2009; da Silva et al., 2009; Guarino et al., 2017). While higher BMI may yield increased energy expenditure necessary for body weight, it could be that the activation of the sympathetic ANS may be an initial or even primary driving force in weight maintenance regulation, possibility advanced in the past (Molfino et al., 2009). In addition, with links between high BMI and baroreceptor dysfunction, hypertension, organ damage, and cardiovascular disease, there may be a potential for therapeutics advanced by sympathetic inhibition.

Our data also show ANS responses that varied with circadian rhythm. Sympathetic responses were both longer in duration and larger in amplitude during afternoon testing. For the parasympathetic features, longer duration responses occurred in afternoon testing, while larger amplitude responses were observed in morning testing sessions. Most past studies relating diurnal variations of autonomic function compare sympathetic and parasympathetic markers between day and night, and have demonstrated, especially in hypertensive patients, sympathetic activity spikes present in early morning (6:00 AM to 8:00 AM), with little change for the rest of the day and nighttime (Panza et al., 1991; Middlekauff & Sontz, 1995; Marfella et al., 2003; Kario et al., 2004). Although we did not test night time conditions, our data suggest systematic variability cycles within the typical 24-hour period, and as such, may offer therapeutic opportunities with novel temporal characteristics.

Past studies of the ANS report heart rate and blood pressure changes in response to stressor tests (Ziegler et al., 1992; Van de Borne et al., 1994; Pagani & Lucini, 2001; Taylor et al., 2003; McGuire et al., 2005; Freeman, 2006; Borresen & Lambert, 2008; Martinez et al., 2010; Thayer et al., 2010; Castiglioni et al., 2011; Rossi et al. 2015; Bellenger et al., 2016; Radtke et al., 2016; Gonzaga et al. 2017; Michael et al., 2017), but there is a dearth of studies that measure many of the remaining signals simultaneously and during controlled autonomic perturbations. The approach to ANS measurement taken in this work shows that parallel expansion of the modalities of raw physiological signals measured broadens the analysis from simple vitals to numerous measures that include temporal measures of HRV, pupillometry, respiration, and EDA.

This initial application of signal processing and machine learning on a set of standard physiological measures presents some challenges. First, this work depended only on cardiovascular changes during all autonomic tests. Other recordings, especially pupil diameter and EDA, were measured and analyzed in preliminary studies not discussed, but the peak average response did not meet the rigorous threshold of 1σ over baseline.

Additionally, other autonomic measurements like photoplethysmogram (PPG), seismocardiogram (SCG), respiratory effort (RSP), pre-ejection period (PEP) of the heart, and other time-domain and frequency-domain heart rate variability measures were not recorded. The reliability of our HRV measure, therefore, suggest that recording and analyzing these other responses might increase the reliability in defining a normal ANS index.

While the focus on a healthy cohort between the ages of 18 and 45 and BMI less than 30 provided an important baseline and the significant differences suggest plausibility for future recording efforts, the sample size was too restricted. Specifically, significant responses were not always measured for this healthy population in sympathetic or parasympathetic duration or amplitude scales, and there may be observable changes in patient populations or individuals outside of these age and BMI constraints. The limitations of our sample extend to the limitation of the recording modalities collected but not analyzed. For example, an impaired EDA has been linked to early stages of diabetic neuropathy (Petrofsky & McLellen, 2009; Khalfallah et al. 2010), and impaired heart rate, EDA, and pupil dilation have been linked with post-traumatic stress disorder (PTSD) (Pole, 2007; Mckinnon et al., 2020), responses that would not have been observed here. Nevertheless, the methods that focus on a healthy cohort and cardiovascular signals generated a reasonable set of healthy responses that may be used to construct a normal template. Such a range of normal responses might well be used to quantify autonomic variations in actual patient populations.

A third limitation is that this method relies on phasic response to a specific battery of autonomic tests, as opposed to continuous changes in recording modalities. Because the method is not ongoing, it cannot provide a continuously updating biomarker to estimate ANS function and balance. Additionally, this battery focuses on cardiovascular responses due to peripheral signaling in the body, and may explain the lack of robust and significant responses in EDA and pupil diameter. Future work may also include other tests, such as a cognitive aptitude assessment, social stress test or mental arithmetic (Tornatzky & Miczek, 1993; Duschek et al., 2009; Bauerly et al., 2019; Gurel et al., 2020), that can induce more central nervous system mediated responses and, therefore, increase the utility of EDA, pupil diameter, and other recorded modalities. Recent work has also shown that biomarkers, like heart rate and PPG amplitude, can be used to predict responses to transcutaneous cervical vagus nerve stimulation (tcVNS) and model dynamic characteristics of an adapting ANS (Gazi et al., 2020). While it is unclear how this may scale for other conditions or interventions, modeling biomarker responses can be applied to continuously monitoring vital-sensing devices to calculate real-time risk scores and further comprehensive index values related to autoimmune health

This method was developed toward creating data-driven approaches to comprehensively and objectively quantify the ANS. Modern methods of computational science, including machine learning and artificial intelligence techniques, have been used to decode complex clinical and experimental data by detecting patterns, classifying signals, and extracting information to inform diagnostic and treatment actions (Wiens & Shenoy, 2018; Norgeot et al., 2019; Debnath et al. 2020). Continuous data from many sensors, including those in this study and adding electroencephalography (EEG) or other neural recording devices, can be used to train such a model on recordings from healthy, able-bodied individuals to characterize ANS balance. Since the battery of autonomic tests can be completed within 30 minutes and all sensors are non-invasively placed, clinical translation can be simple. Future studies could focus on varying patient populations, as disturbances in autonomic regulation have been described in a variety of diseases, including those resulting from focal injury, such as spinal cord injuries (Krassioukov et al., 2012) and stroke (Dütsch et al., 2007), and diffuse disorders, such as sepsis and infection (Badke et al., 2018; Ferreira & Bissell, 2018), rheumatoid arthritis (Koopman et al., 2016; Koopman et al., 2017), Crohn’s disease (Engel et al., 2015), and diabetes mellitus (Verrotti et al., 2014; Serhiyenko & Serhiyenko, 2018). Additionally, dysautonomias have been described in numerous cardiovascular conditions (Broadstone et al., 1991; Kishi, 2012; Carthy, 2013; Vinik et al., 2013; Shen & Zipes, 2014) and central nervous system disorders, including Alzheimer’s disease (Femminella et al., 2014), Parkinson’s disease (Goldstein, 2014), Huntington’s disease (Kobal et al., 2010; Diago et al., 2017), and psychiatric conditions including depression, schizophrenia, and PTSD (Jung et al., 2019), among others. Another possible application could be evaluation of targeted neuromodulation; specifically, there is clinical interest in stimulating the vagus nerve, which is involved with responses in cardiovascular, pulmonary, gastrointestinal, renal, hepatic, and endocrine systems (Chavan et al., 2017; Pavlov et al., 2018). Vagus nerve stimulation (VNS) has been used in previous studies for multiple conditions, including refractory epilepsy (Stefan et al., 2012; Rong et al., 2014), depression (Rong et al., 2016; Kong et al., 2018), PTSD (Bremner et al., 2020), pre-diabetes (Huang et al., 2014), tinnitus (Shim et al., 2015), stroke (Redgrave et al., 2018), and others, including oromotor dysfunction, rheumatoid arthritis, and obesity (Guiraud, et al., 2016). These studies have used a range of electrical stimulation settings and sites, and there is no optimal dose or set of parameters (Badran et al., 2018); the mechanism of VNS and responses are not well understood. Studies that report significant effects of VNS on HRV, pupil diameter and evoked potentials are mixed or report no significant changes but the preliminary effects on clinical populations are clear (Libbus et al., 2017; Burger et al., 2020; Gurel et al., 2020). A set of biomarkers or calculated features to accurately and consistently quantify the ANS in a patient-centered approach can be extremely helpful in a number of clinical applications.

The sympathetic and parasympathetic parameters determined in this study will be valuable to diagnose autonomic function and underlying disorders, as well as predict responses to targeted modulated therapies. While the use of autonomic modulation has shown promise in treating cardiovascular, autoimmune, and nervous system disorders, the template matching method in this work can offer additional insight towards the effects of stimulation and medication in patient populations.

## ADDITIONAL INFORMATION

### Data Availability

The data that support the findings of this study are available from the corresponding author upon reasonable request.

### Author Contributions

SD, TL, and TPZ conceptualized and designed the study. SD and TL acquired and analyzed all data. MB, RMS, DPB, SZ, and BTV contributed to interpretation and discussions at multiple stages of manuscript development. SD, TL, MB, RMS, DPB, SZ, BTV, and TPZ contributed to drafts of the manuscript. All authors critically revised and approved the final version of the manuscript and take responsibility for the integrity of the work. All persons designated as authors qualify for authorship, and all those who qualify for authorship are listed.

## Acknowledgment

The authors would like to thank all participants who took part in testing sessions.

